# Further genetic diversification in multiple tumors and an evolutionary perspective on therapeutics

**DOI:** 10.1101/025429

**Authors:** Yong Tao, Zheng Hu, Shaoping Ling, Shiou-Hwie Yeh, Weiwei Zhai, Ke Chen, Chunyan Li, Yu Wang, Kaile Wang, Hurng-Yi Wang, Eric A. Hungate, Kenan Onel, Jiang Liu, Changqing Zeng, Richard R. Hudson, Pei-Jer Chen, Xuemei Lu, Chung-I Wu

## Abstract

The genetic diversity within a single tumor can be extremely large, possibly with mutations at all coding sites (Ling et al. 2015). In this study, we analyzed 12 cases of multiple hepatocellular carcinoma (HCC) tumors by sequencing and genotyping several samples from each case. In 10 cases, tumors are clonally related by a process of cell migration and colonization. They permit a detailed analysis of the evolutionary forces (mutation, migration, drift and natural selection) that influence the genetic diversity both within and between tumors. In 23 inter-tumor comparisons, the descendant tumor usually shows a higher growth rate than the parent tumor. In contrast, neutral diversity dominates within-tumor observations such that adaptively growing clones are rarely found. The apparent adaptive evolution between tumors can be explained by the inherent bias for detecting larger tumors that have a growth advantage. Beyond these tumors are a far larger number of clones which, growing at a neutral rate and too small to see, can nevertheless be verified by molecular means. Given that the estimated genetic diversity is often very large, therapeutic strategies need to take into account the pre-existence of many drug-resistance mutations. Importantly, these mutations are expected to be in the very low frequency range in the primary tumors (and become frequent in the relapses, as is indeed reported (1-3). In conclusion, tumors may often harbor a very large number of mutations in the very low frequency range. This duality provides both a challenge and an opportunity for designing strategies against drug resistance (4-8).

**One Sentence Summary:** The total genetic diversity across all tumors of a single patient, with large number of low frequency mutations driven by neutral and adaptive forces, presents both a challenge and an opportunity for new cancer therapeutics.

## Introduction

Recent genomic sequencing has led to the increasing identification of diverging genetic clones within a given tumor (1, 9-24). The term “tumor” is used here to designate a population of cancerous cells that occupy a discrete space. Theories on migration and population differentiation may therefore be applicable to the evolution of multiple tumors (25, 26). In a single HCC tumor, Ling et al. (2015) have estimated the genetic diversity to be extremely large with millions of coding region mutations. The analysis is extended to multiple tumors in this study. Since the total genetic diversity in all tumors of the same individual may likely determine the outcome of treatments, dispersed tumors are particularly challenging.

Genetic diversity in tumors is governed by the evolutionary forces of mutation, genetic drift and selection (25-28). For multiple tumors, cell migration would be an additional force. While selection is often assumed to drive some clones to grow faster than others (20, 24), it is in fact the only force that may or may not be in operation. Its action needs to be proven, rather than assumed. Here, we aim to test the action of selection both within and between tumors by comparing the observations with the null model, in which all clones have the same neutral growth rate. Regardless of whether and how selection influences the genetic diversity, a large amount of neutral diversity, possibly orders of magnitude higher than the “adaptive” diversity, is expected to exist. Such neutral diversity may likely have biological consequences when the environment changes, a phenomenon discussed in the companion study (see Ling et al. 2015 on the Dykhuizen and Hartl effect). The level of neutral diversity can be computed (29) but has not been systematically formulated and measured.

Multiple tumors are evolutionarily more complex than single tumors as they would encounter divergent micro-environments and experience cell migration. No less important is an observational bias due to cell motility. After cancerous cells migrate and proliferate, tumors of a larger size due to a growth advantage would be preferentially detected, whereas newly seeded tumors that grow neutrally would often be too small to see. As a consequence of this observational bias, the role of selection would appear much larger between tumors than within tumors. Even though neutrally growing tumors may outnumber adaptively growing ones by orders of magnitude, the latter would still be predominantly observed and sampled. In contrast, small clones could be sampled without bias within the same tumor, whose boundary is delineated (for example, Fig. 1A of Ling et al. 2015). The bias is illustrated in Fig. 1A.

**Fig. 1.**
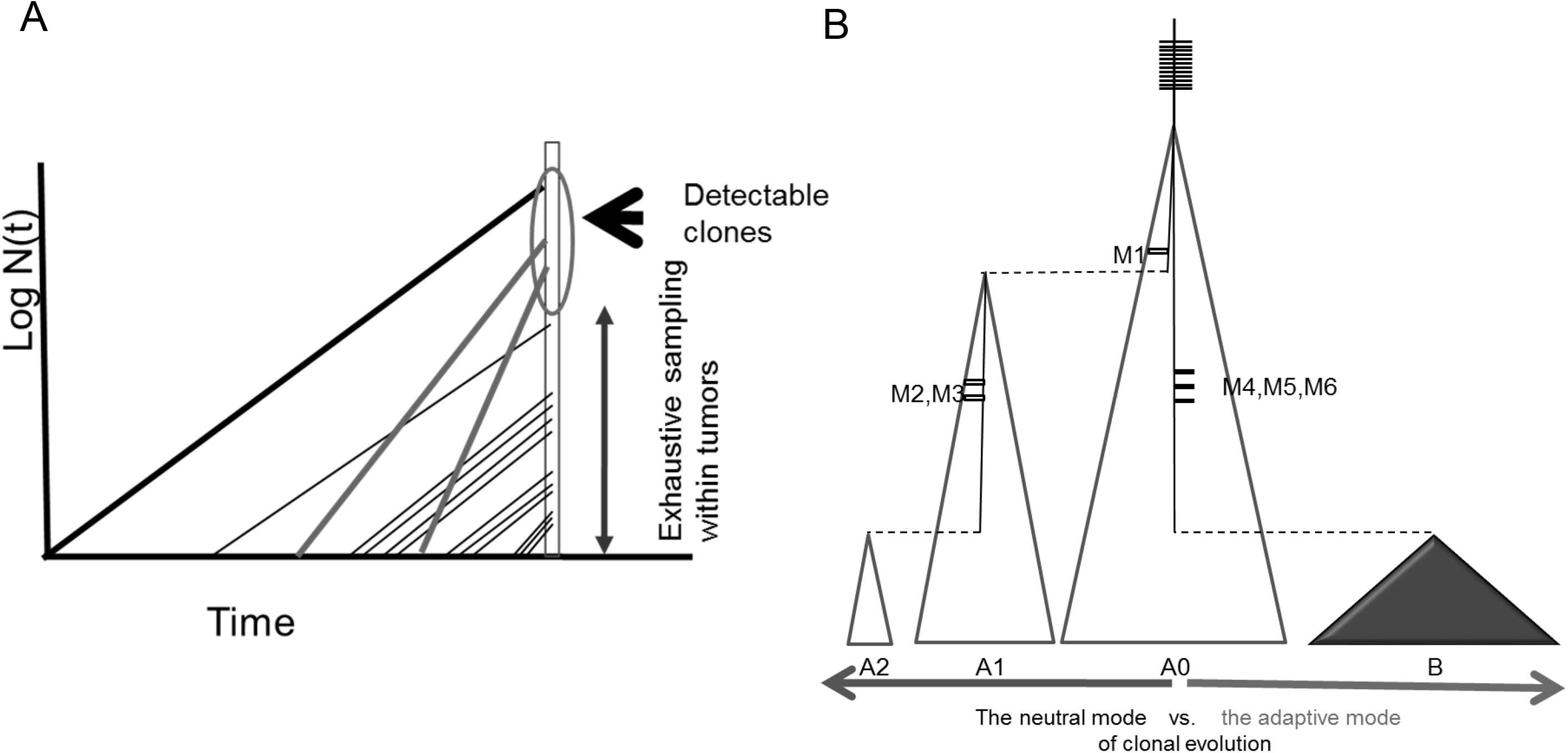
Models of tumor evolution. (A) Clonal growth and observational biases – Exponentially growing clones are depicted as straight lines with parallels indicating clones of equal growth rate. New clones emerge more frequently as the population expands. There is an inherent bias in the detection of multi-tumor cases because only large clones are observable by size, marked by the oval. Larger clones usually have a growth advantage as shown by the red lines. In a single tumor, smaller clones can still be sampled systematically within the circumscribed boundary. (B) Illustration of neutral vs. adaptive growth of clones – A0 is the ancestral clone, the width reflecting the clone size on a logarithmic scale. The dashes on the trunk and branches are somatic mutations (M). A1 and A2 have the same growth rate as A0 and are smaller due to their late onsets of growth, indicated by the acquisition of additional mutations. A0 – A2 are hence neutral-growth clones. In contrast, clone B starts to grow much later but becomes larger in a shorter time. Clone B has a growth advantage over the others.

The comparison of the evolutionary forces within-and between-tumors will be informative in two ways. First, it will provide insights into how natural selection influences the level and pattern of genetic diversity in tumors. Second, it will permit an account of the total diversity in multi-tumor cases by providing estimates of the unobserved tumor mass. In this study, we carry out whole-exome sequencing on 12 cases of multi-tumor hepatocellular carcinomas (HCCs) (see Table 1 and Table S1). Exome sequencing is used on 1 – 6 tumor samples per individual and another sample from the non-tumor tissue of the same liver (see Supplementary Materials (SM) section A1, Table S1 and S2). In total, 74 cancerous samples were sequenced and/or genotyped.

**Table 1.** Summary of cell clones in the 12 cases of multi-tumor HCCs.

| Case | # of Tumors | # of samples from tumors<sup>a</sup> | # of founder Mutations<sup>b</sup> | # of cell clones | # of adaptive clones<sup>c</sup> | Diversity between tumors | Diversity within tumors |
| --- | --- | --- | --- | --- | --- | --- | --- |
| **Overall low diversity** |  |  |  |  |  |  |  |
| HCC-9<sup>d</sup> | --<sup>d</sup> | 5 (2) | 100 | 2 | 0 | little diversity (possibly high migration rate) |  |
| **Neutral diversity only** |  |  |  |  |  |  |  |
| HCC-6 | 2 | 6 (2) | 50, 31 | 3 | 0 | independent origin | neutral |
| **Neutral diversity within tumors and/or adaptive diversity between-tumors** |  |  |  |  |  |  |  |
| HCC-8 | 3 | 5 (2) | 39 | 4 | 3 | adaptive | NA |
| HCC-7 | 3 | 6 (2) | 49 | 5 | 3 | adaptive | neutral |
| HCC-12 | 3 | 12 (3) | 33 | 5 | 3 | adaptive | neutral |
| HCC-11 | >10 | 10 (6) | 30 | 11 | 8 | adaptive | NA |
| **Adaptive diversity within- and between-tumors (within-tumor diversity transient; see Fig. 4B)** |  |  |  |  |  |  |  |
| HCC-1 | 3 | 9 (3) | 21 | 5 | 3 | adaptive | Adaptive, transient |
| HCC-3 | 3 | 7 (2) | 41 | 2 | 1 | adaptive | Adaptive, transient |
| HCC-4 | 7 | 7 (1) | 58 | 3 | 2 | adaptive | Adaptive, transient |
| **Uninformative cases** |  |  |  |  |  |  |  |
| HCC-5 | 2 | 2 (2) | - | 2 | - | independent origin | NA |
| HCC-2 | 2 | 3 (1) | 12 | 2 | 0 | NA | NA |
| HCC-10 | 2 | 2 (1) | 30 | 1 | - | NA | NA |

<sup>a</sup> Total number of samples genotyped/sequenced (the number of samples sequenced).
<sup>b</sup> Founder mutations are those shared by all cell lineages.
<sup>c</sup> A case with only one lineage would have 0 adaptive lineages.
<sup>d</sup> HCC-9 is a cluster of >> 20 small nodules (see Fig. 2A).

## Results

### I. Theory

#### Theory of the size distribution of tumors with cell migration

To understand the forces driving multi-tumor evolution, it is necessary to derive the size distribution of clones. Within the same tumor, clones are defined by new mutations but, in multi-tumor cases, a new tumor seeded by migrant cells can be considered a new clone as well. By substituting the rate of migration for the mutation rate, we can derive the size distribution of tumors, just as we calculated clone sizes in Fig. 4 of Ling et al. (2015), using the infinite-allele model (25, 26, 30, 31). A cell clone is thus either a population of cells sharing the same mutation (a mutation clone) or cells originating after the same migration event (a migrant clone, or a new tumor).

In formulating the clone size distribution, the neutral null model assumes that cells divide and mutate (or migrate) with a given probability without invoking selection or other complex interactions (Ling et al. 2015). Let the migration rate (number of migrant cells per cell per generation) be m. Given any m, we ask i) how many tumors are visible, defined as those that are larger than, for example, 1% of the total tumor mass, and ii) how many are too small (< 1%) to see? Various tumor migration models (32-35) make similar predictions on visible tumors and the dynamic model of Iwata et al. (32) is adopted here.

The model assumes that a tumor grows exponentially with a rate of r (29). The number of emigrant cells at time t is m[N(t)]^α^ = me^αrt^, where α is a fractal dimension factor that corrects the spatial effect on cell migration. When only cells on the periphery of a three-dimensional sphere are capable of emigration, α = 2/3. Let G(x) be the number of migrant clones which have more than x cells at time T. For neutrally growing tumors (see SM section B3),

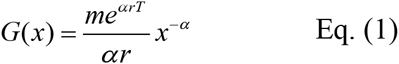

Eq. (1) has been extended to include growth advantages. In the extension, the newly seeded tumor has a growth rate, s, which may be higher or lower than r (the growth rate of the parent tumor). We assume that s follows an exponential distribution with the mean of r. With this distribution, the growth rate of most new tumors would be lower than the parental tumor, but a few would have a much greater growth rate. G’(x) is the

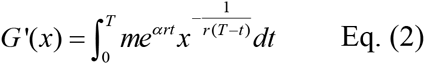

Both G(x) and G’(x) (see SM section B3 for the derivation) will be used to estimate the total tumor mass that will include the unobserved clones.

#### Criteria for determining neutral vs. adaptive growth of cell clones

In the accompanying study of clone size distribution, the clones *collectively* do not deviate from the null model (Ling et al. 2015). We now investigate the growth dynamics of each clone individually in order to capture rare adaptive events amidst the neutral pattern. A clone is suggested to have a growth advantage over another if the former has grown larger in a shorter time. The size of cell clones can often be delineated for migrant clones that form discrete and separate tumors. The age of a clone is determined by mutation accumulation. In this study, only the parent - descendant pairs are used to determine the relative age (older vs. younger). In the illustration of Fig. 1B, A1 clone and its parent A0 share all the founder mutations but A1 has an extra mutation, M1, which emerges in a cell within the A0 clone. A1 thus starts its expansion later than A0. In this figure, cell lineages are drawn as triangles to denote their expansions from a single cell. A0, A1 and A2 follow the neutral growth dynamics by which cell clones that emerge later grow to be smaller (see legends). In contrast, clone B starts to grow late, after its progenitor cell acquires mutations M4 – M6. Its large size thus suggests a growth advantage over A0.

The relative sizes of clones can be complemented by another measure of growth advantage embedded in the sequence data. This measure is the average number of mutations per cell, *L*, that have accumulated since the origin of the clone. Its expression,

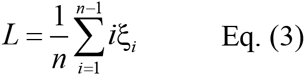

was first proposed by Zeng et al. (36) who denote ξ_*i*_ as the number of sites where the mutant is found in i of n sequencing reads (see SM section B1). By the measure of *L*, a clone is said to have a higher fitness than another if it has accumulated more mutations, likely through more cell divisions, in a shorter time.

### II. Observations on the diversity between tumors

The cases of HCCs used in this study are summarized in Table 1. Two of the 12 cases are of independent origin (HCC-5 and -6) because tumors from the same patient share no somatic mutations. (When tumors do share somatic mutations, >60% of these mutations are shared.) Independently originated tumors, interesting as they are, offer no information on the evolution of inter-tumor diversity (see Table 1).

#### Neutral clonal growth between distinct tumors

Among the 9 informative cases, HCC-9 is exceptional in having dozens of small tumors clustered together and giving a diffuse appearance (Fig. 2A). Samples from 5 tumors were taken, of which 2 were sequenced and all 5 were genotyped. These samples are nearly identical, sharing all 96 somatic mutations (see Table S3) and suggesting comparable ages of these tumors. HCC-9 exhibits a neutral pattern in which clones of a similar age are similar in size (Fig. 1B) with no evidence of growth differences among tumors. Such a neutral pattern suggests a high rate of cell migration, to be discussed below.

**Fig. 2.**
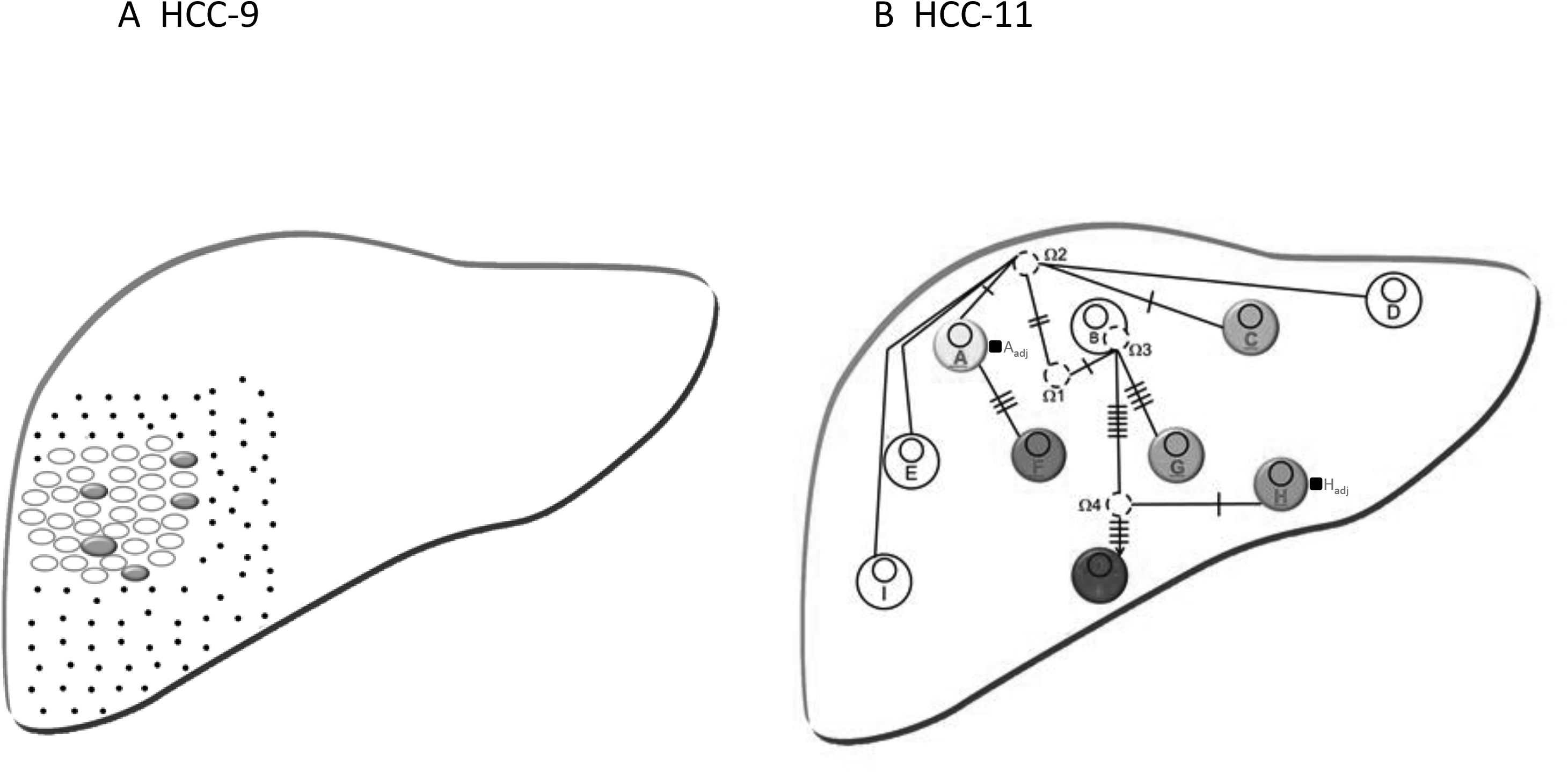
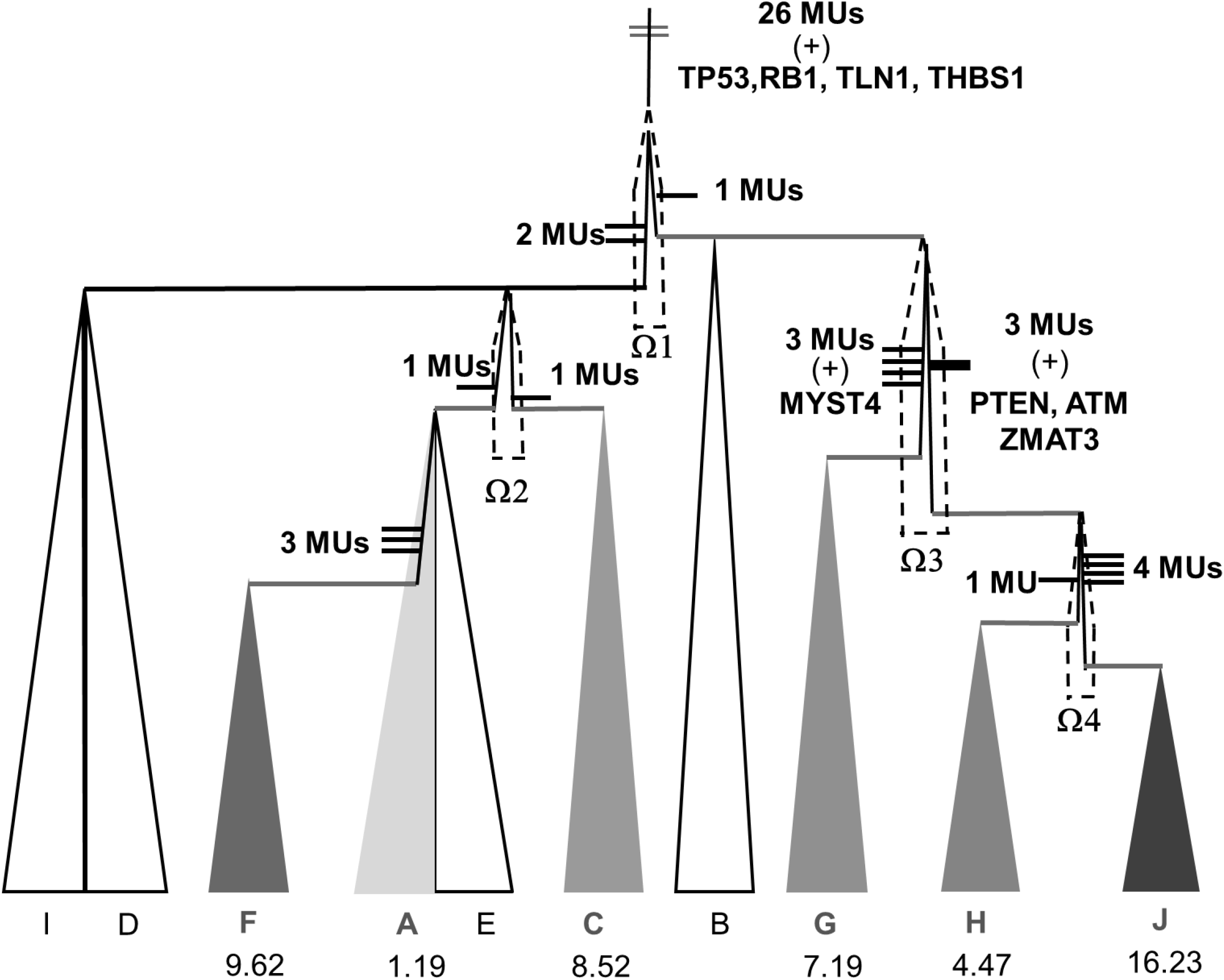
Two cases with large numbers of tumors. (A) HCC-9 is an unusual case of many small tumors with a “diffuse” appearance. Dots suggest many small tumors on the fringe contributing to this appearance (see Fig. 3C for simulations). All five samples share 96 coding mutations and only one sample has an extra mutation. By the criteria of Fig. 1, this is the only case of neutral growth between tumors. (B) In HCC-11, distinct tumors are denoted nodules A to I, 6 of which are sequenced and all are genotyped. Ancestral tumors can be inferred and are labeled Ω’s. Nonsynonymous mutations linking tumors are shown as ticks. A_adj_ and H_adj_ are the non-tumor samples adjacent to tumor A and H. (C) The genealogy of the 10 sampled tumors. Note that the descendant clone, although starting to grow later, grows to be larger than the ancestral tumor in the mode of clone A0 → clone B of Fig. 1C. The sampled tumor may originate in an ancestral clone that is too small to detect. Genes with potential cancer-driving mutations are given and MU’s denotes other nonsynonymous mutations. The *L* value (Eq. 3), given below each sequenced tumor, should reflect the activity of cell divisions of that tumor. Sequenced samples are shown with filled color. The triangle indicates the growth of a cell lineage.

#### Adaptive clonal growth between distinct tumors

Multiple tumors growing neutrally, as seen in HCC-9, are not common. In general, new tumors need to have a growth advantage to attain a mass sufficient for detection. A dramatic example is HCC-11; samples from its 10 tumors were taken (Fig.2B), of which 6 were sequenced and all 10 were genotyped. In total, 11 cell lineages are observed or inferred as shown in Fig. 2C. Overall, 30 coding mutations are shared by all lineages, but only 1-4 mutations are associated with the emergence of each lineage. The spatial pattern of divergence and the inferred migration routes show neighboring tumors to be genealogically closer than distant ones.

As shown in Fig. 2C, 8 separate events of accelerated clonal growth can be inferred from the 10 different samples of HCC-11. In these events, derived lineages have grown to be larger in size (N) or have accumulated more mutations (larger L) than the parental ones, many of which are inferred and labeled Ω in Fig. 2C. Three of the 4 Ω lineages have a genotype not represented by any of the samples and are probably too rare to be sampled. The 6 sequenced tumors (solid triangles) are larger than the ancestral clones in size but started to grow later. Interestingly, the oldest clone A has the smallest *L* value while the youngest clone, J, has the largest L, suggesting that the younger clones have divided more often in a shorter time period. The spatial pattern of these cell clones and the number of mutations separating them are depicted in Fig. 2C. Six other cases, HCC-1, HCC-3, HCC-4, HCC-7, HCC-8 and HCC-12 are similar to HCC-11 in their support of adaptive divergence between tumors. All of these cases have 2-3 tumors except HCC-4, which has 7. HCC-7 will be analyzed later for intra-tumor differentiation, while the other cases are shown in Fig. S1 – S4.

Studies in cancer genomics often aim to identify tumor-driving mutations. However, the identities and functions of the drivers are usually obscured by the much larger number of passenger mutations (37, 38). For this reason, the comparison between two cancerous clones might be particularly informative (12, 23) as long as the clones being compared are known to be adaptively divergent (see Fig. 1B and Fig. 2C). When two such clones differ by only a few mutations, as is commonly observed, the identification of drivers would be simpler.

In this study, we identified 650 somatic mutations in 594 genes associated with the 12 HCC cases (Table S3). Most of these mutations are “fixed” mutations found in all samples from the same individual, as defined in Ling et al. (2015). Among the 76 “polymorphic” mutations that were found in some but not all tumor samples, 17 are of greater interest than the rest (see Table 2). Ten of them are the sole nonsynonymous mutation between two clones, one of which is derived from the other and have a growth advantage. The remaining 7 are in a small set of 2-3 mutations which also define such a parent-descendant expansion. These 17 mutations are putative drivers because no other coding mutations were found.

**Table 2.** Nonsynonymous mutations in the exome of adaptively-evolving tumors, relative to the parent tumors. Only those clonal relationships with 1 to 3 mutations are shown.

| Case | Clonal relationship<sup>a</sup> |  |  | Gene name |
| --- | --- | --- | --- | --- |
| **HCC-1** | $\Omega$ | → | R2 | CCNG1 |
| | T1 | → | R1 | P62 |
| **HCC-3** | T1-4 | → | T1-1 | ATP8A2, KIAA0556 |
| **HCC-4** | T2 | → | T3 | TNFRSF10D |
| **HCC-7** | T1-3 | → | T1-1 | LRRTM3 |
| **HCC-8** | $\Omega$ | → | T1 | IL25, ITIH5, HLA-DQB1 |
| **HCC-11** | Ω1 | → | Ω2 | CAPSL, F7 |
| | Ω1 | → | B | CUBN |
| | Ω3 | → | H | FUT8 |
| | Ω2 | → | A | EFS |
| | Ω2 | → | C | PKHD1 |
| **HCC-12** | Ω2 | → | T0 | MKI67 |
| | Ω1 | → | Ω2 | IGFL4 |

<sup>a</sup> A → B indicates that B, a clone growing out from clone A, has a growth advantage over A as illustrated in Fig. 1B (e.g. A0 → B). See Fig. 2, Fig. 5 and Supplementary Fig. S1 - S4 for clonal designations.

Table 1 therefore presents an alternative means for inferring driver mutations, in complement to the conventional approach which relies on the sharing of mutated genes among tumor cases (37, 38). However, drivers identified by the two approaches may differ in many ways. For example, the sequence of mutation accumulation leading to a growth advantage is defined in Table 1 but not in the conventional method. Hence, studies that combine and/or compare the two approaches will be most interesting.

#### Total number of tumors – observed and unobserved

The presence of multiple clonally related tumors would suggest the existence of undetected tumors since cell migration may be a continual process. The detection of circulating tumor cells lends support to that inference (39). Given the number of observed tumors, models can be constructed to inform about the number of the unobserved, which can then be validated experimentally. Using Eq. (1) and (2), we model the relative numbers of observed and unobserved tumors as shown in Fig. 3A and 3B. For a first approximation, 1% of the total tumor mass is assumed to be the dividing line between the observed and the unobserved.

**Fig. 3.**
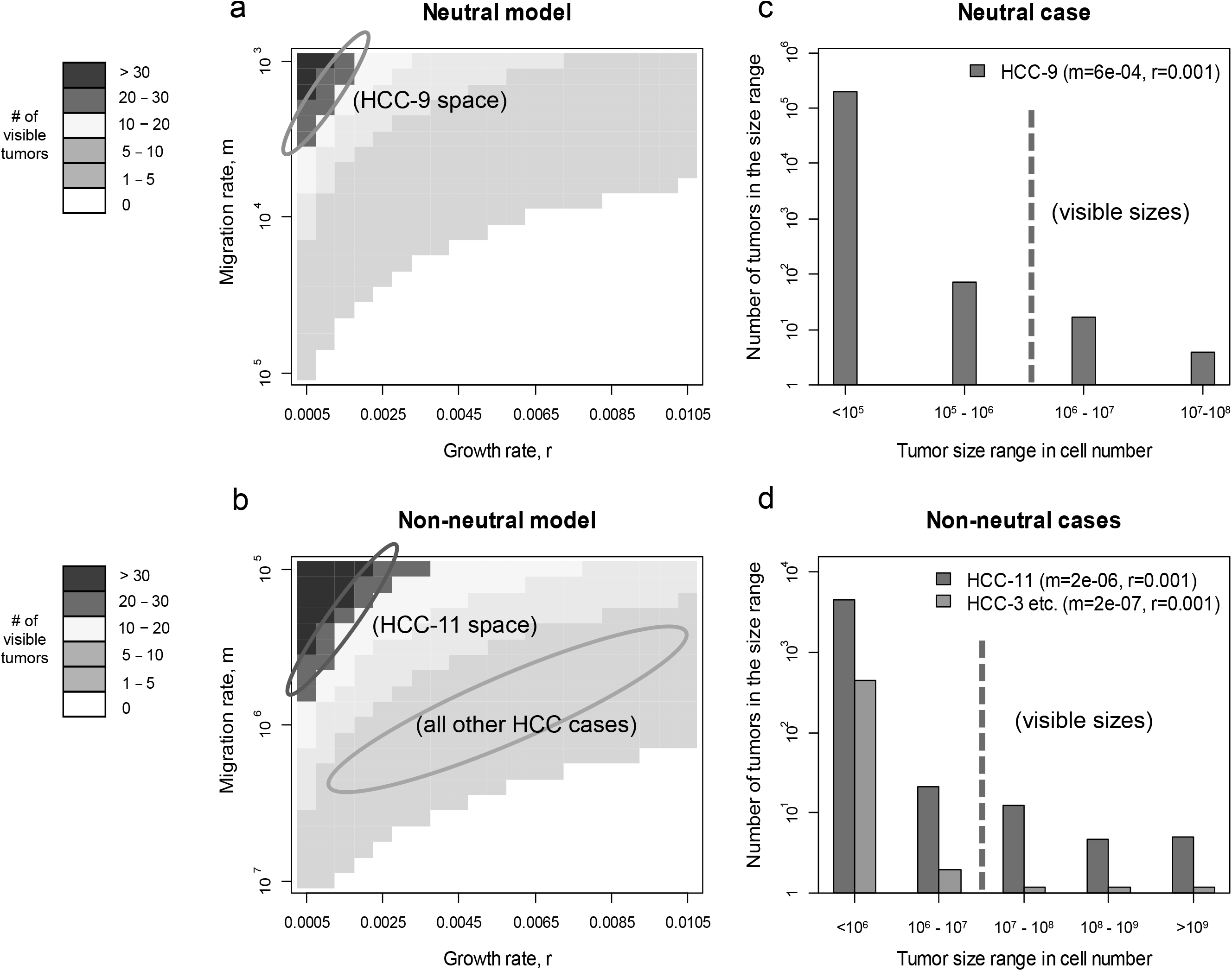
The expected numbers of visible tumors (A – B, in heatmaps) and the size distributions of all tumors (C - D). Visible tumors are defined as those that are larger than 1% of the total tumor mass, which is estimated to be 10^8^−10^9^ cells for the HCC cases. (A, C) Neutral growth model and HCC-9. (B, D) Adaptive growth model and all other cases. The heatmaps in the left panels show the number of visible tumors under different growth rates (r) and migration rates (m). The ovals mark the possible parameter space applicable to the HCC cases. The bar plots in the right panels show the size distributions of tumors for the corresponding cases. The red broken lines delineate visible and non-visible tumors.

As defined by Eqs (1-2), the total mass of tumor is a function of r (the growth rate) while the number of tumors is a function of m (the cell migration rate). Given the same r, there would be more, but smaller, tumors as m increases (see Fig. 3A). Most importantly, the effect of migration on tumor evolution depends greatly on the presence of selection. The comparison between the neutral and adaptive growth shows this dependence clearly. As the migration rate decreases continuously from the top of Fig. 3A to the bottom of Fig. 3B, the calculations suggest that the neutral growth mode would require 100 fold more migration to yield a comparable number of detectable tumors.

HCC-9 has >20 tumors, most of which appear to have the same neutral growth rate, as suggested above. It is possible to see so many neutrally growing tumors only if the migration rate is very high and the growth rate is low (the upper left corner of Fig. 3A). The seeding of this many tumors of equal size had to occur when the primary tumor was small, which would require a high rate of cell migration (m > 0.3×10^−3^ in Fig. 3A). At such a high migration rate, the estimated number of unobserved tumors should be large. Fig. 3C indeed predicts > 50 tumors in the 1% size range of the total cell mass, as well as many smaller ones. The pathological report of “diffuse tumors” can thus be explained by the presence of numerous tiny tumors.

In all other cases, growth advantage drives the evolution of multiple tumors, allowing m to be much lower than under the neutral model (Fig. 3B). Between Fig 3A and 3B, m is different by two orders of magnitude but yields comparable numbers of visible tumors. HCC-11 has > 10 visible tumors and most other cases have fewer than 5 (Table 1). The suitable ranges for m and r that cover the observations are marked in Fig. 3D. Given the r and m values, the size distributions of the tumors are shown in Fig. 3D. We estimate that HCC-11 and the remaining cases have more than 1,000 and 100 unobserved tumors, respectively.

#### Evidence for tumors below the observable range

If migrant cancer cells indeed form tumors that are too small to see, it may be possible to detect them in non-tumor samples by molecular means. Furthermore, cases with higher migration rates should have more cancer cells in non-tumor samples. We analyzed HCC-11 and HCC-1, which have, respectively, the highest and lowest rate of cell migration among the adaptive cases. From HCC-11, three non-tumor samples were taken – A_adj_ (a sample adjacent to tumor A, see Fig. 2B), H_adj_ (adjacent to the most peripheral tumor H) and CB (circulating blood). All 47 coding region mutations found in the tumors were genotyped in these samples (Fig. 4A). As low frequency variants are individually unreliable, it is necessary to compare multiple variants for their overall trend. Non-tumor samples were calibrated against the tumor samples, which have the mutations at a frequency of about 45% (50% being the expectation for heterozygous mutations). The non-tumor samples, A_adj_, H_adj_ and CB, are estimated to have ∼ 4.8%, ∼0.8% and ∼ 10% of the tumor mutations, respectively (Fig. 4B). Since A_adj_ is near the origin of HCC-11 tumors and H_adj_ is at the periphery, the difference is expected. The CB sample has the highest percentage of tumor cell DNA, but much of which may have come from degraded cells. The results support the conjecture that many cancer cells imbue the liver containing HCC-11 without forming large and visible tumors.

**Fig. 4.**
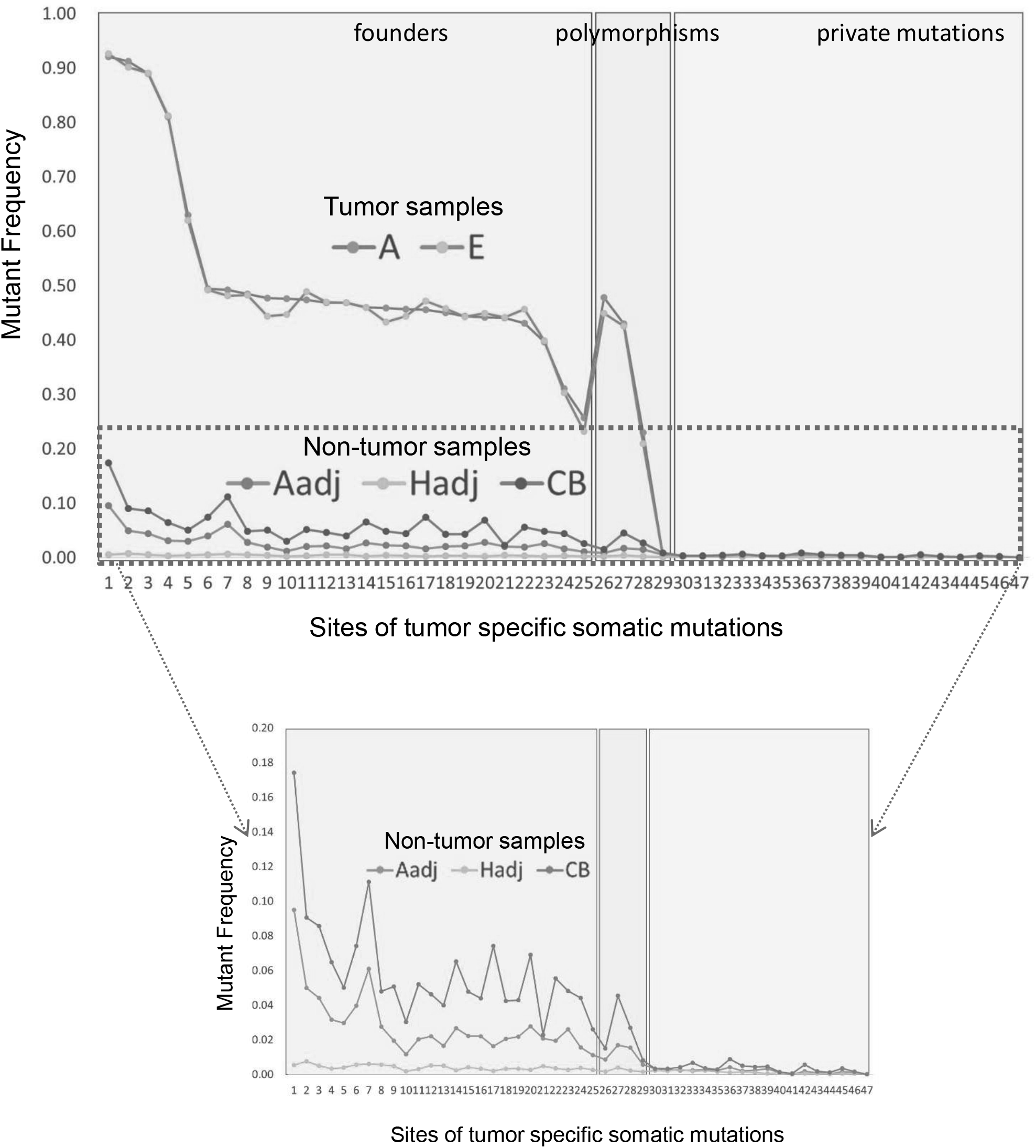
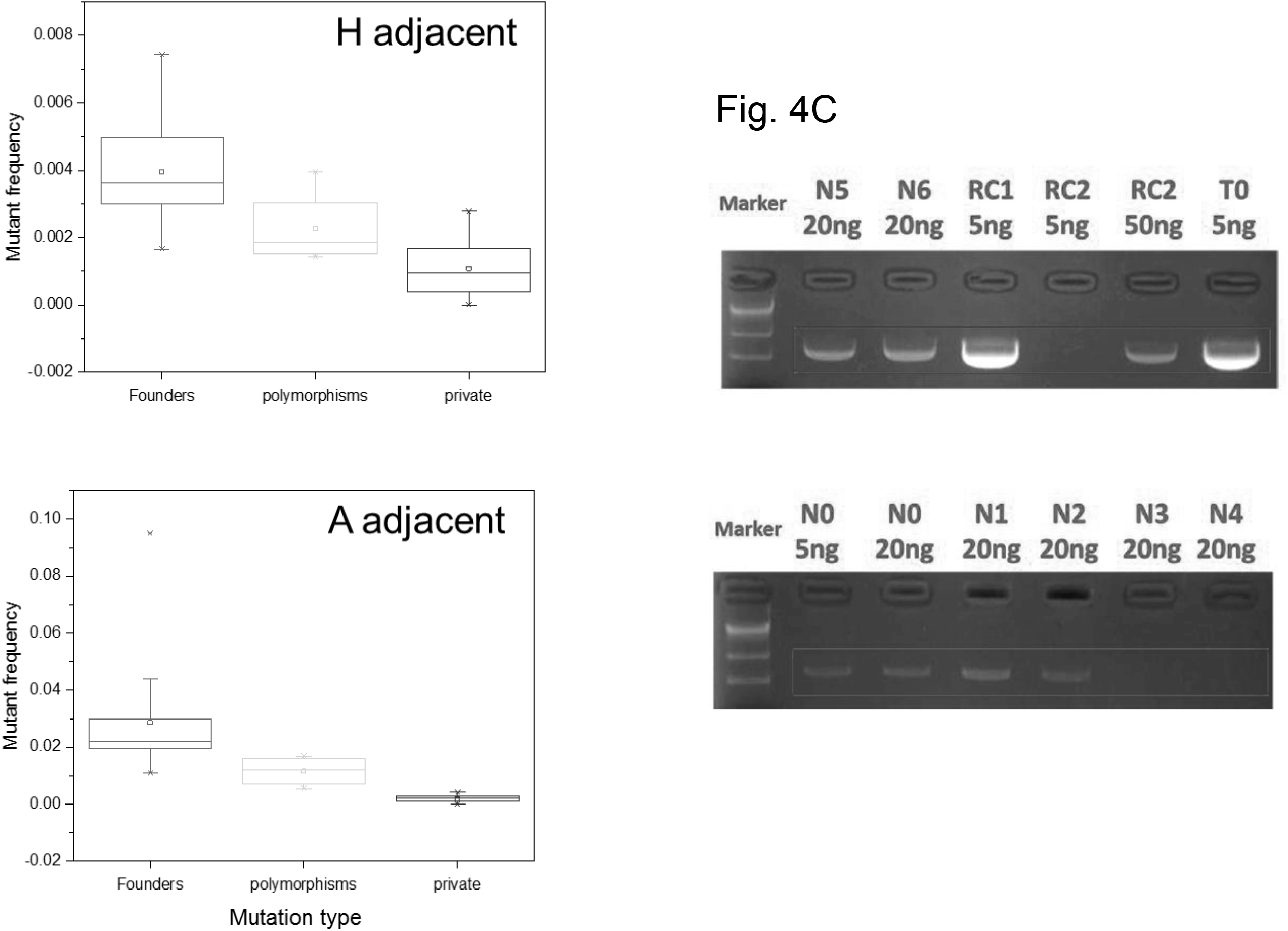
Detection of cancer cells in non-tumor samples. (A) Non-tumor samples (A_adj_, H_adj_, CB; see text and Fig. 2B) of HCC-11 are compared with the tumor samples (nodules A, E) for the frequencies of somatic mutations detected in tumors. The 48 somatic mutations fall into 3 classes – founders (present in all tumors), polymorphic mutations (present in 3-9 tumors) and private mutations (present in 1-2 tumors). Profiles in the non-tumor samples follow the same trend as those in the tumors, but at a much lower level. Frequency profiles have been validated by PCR-NGS sequencing, with an average coverage of > 10,000X. (B) Box plots of the profiles of the A_adj_ and H_adj_ samples of HCC-11 are shown. The founder mutations are most informative due to their high frequencies. The A_adj_ sample is estimated to have more than 4% cancer cells while the H_adj_ sample has 0.7%. (These percentages are twice the median value of the Y-axis values due to the heterozygosity of the mutations.) A_adj_ is near the center of tumor distribution while H_adj_ is at the periphery and the difference suggests distance-dependent cell movement. (C) Non-tumor samples (N1 – N7) of HCC-1 are compared with the tumor samples (T0, RC1 and RC2) for the frequencies of an HCC1-specific mutation. This mutation is a large deletion specific to the T0 and RC1 tumors and is detected by the presence of the junctional sequence across the breakpoint (12). Based on the amount of the template and the PCR results, the relative abundance of the HCC1-specific markers fall into these categories: High (RC1 and T0), medium (N0), rare (N1, N2, N5 and N6), very rare or absent (RC2, N3 and N4).

From HCC-1 (Fig. S1), seven non-tumor samples (N0-N6) were taken. For comparison, HCC-1 should be considered a one-tumor case because two of the three tumors are recurrences, unobserved at the first surgery (12). Single nucleotide mutations from tumors are not detectable in these samples (see (12)), corroborating the analysis that m is relatively low in HCC-1. However, two of the HCC-1 tumors carry a large deletion (Δ5q), the breakpoints of which have been identified (12). PCR products across the breakpoint are specific to the cancer cells, thus making their detection feasible even at very low frequencies. Fig. 4C shows five non-tumor samples with some Δ5q-bearing cells. These five samples in fact have more Δ5q-bearing cancer cells than the RC2 tumor, another tumor seeded early during tumor growth. In this least migratory case, cancer cells can still be found in the majority of non-tumor samples from the liver, albeit at a much lower frequencies than in samples of HCC-11.

### III. Observations on the diversity within tumors

Adaptive differences in growth between tumors seem pervasive although much of it could be due to observational bias (see Fig. 3A vs. 3B). Adaptive differences within tumors (between clones of the same tumor) have also been commonly assumed (20, 24). Since a rigorous test did not support this view in one HCC case (Ling et al. 2015), we now expand the analysis to additional cases (see Table 1).

#### Adaptive diversity in the transition from intra-to inter-tumor variation

In three cases (HCC-1, -3, and -4), the diversity within tumors overlaps with the inter-tumor diversity and is in the transitional phase. The adaptive diversity between tumors must first emerge within tumors, much like inter-specific differences first originate as intra-specific variation (40). Such intra-tumor diversity represents the early stages of the process of inter-tumor divergence. The question is where the adaptive mutation originates - in the old or the new tumor? Fig. 5A shows the scheme in which the diversity originates in the old tumor and Fig. 5B shows it originating in the new tumor. In these schemes, the descendent clone shown in dark red has all the mutations of the parental one, shown in pink. A tumor may have more than one clone and a clone may be present in more than one tumor.

The implications of the two schemes are quite different. In the scheme of Fig. 5A, the divergence between the older and younger tumors originates within the older tumor. It is possible that there is no fitness difference between the two clones and the colonization by the younger clone (dark red) is a matter of chance, similar to the founder effect in natural populations (40). In comparison, Fig. 5B presents a scheme of adaptive evolution in which the ancestral clone (pink) seeds the new tumor. The descendent clone subsequently surpasses the ancestral one to become dominant. To test the two models, the diversities within-and between-tumors must overlap and only three cases (HCC-1, -3, and -4) are informative in this respect.

**Fig. 5.**
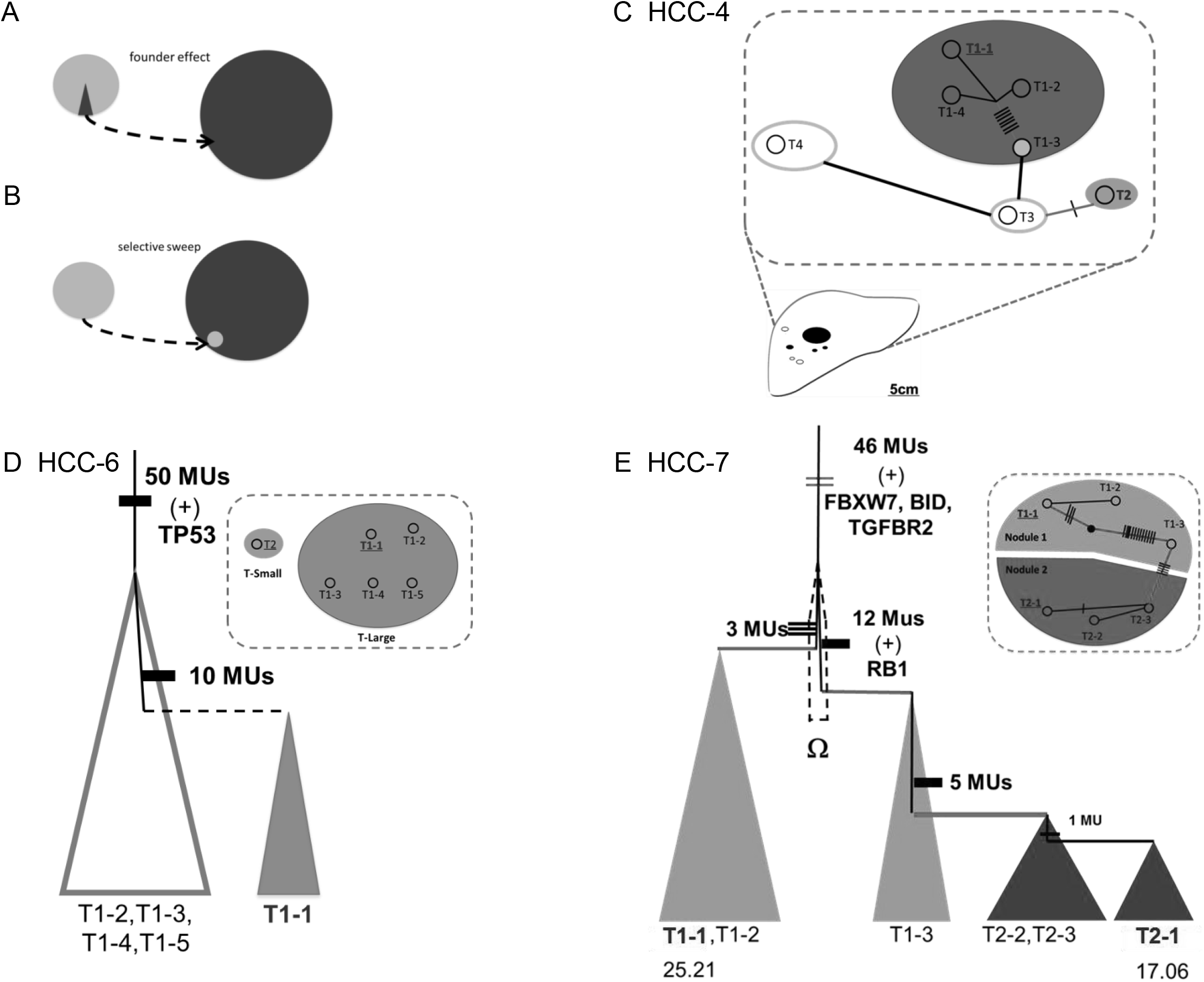
Within-tumor diversity. (A - C) Analysis of within-tumor diversity that is in transition to inter-tumor divergence. There are two models on the origin of inter-tumor adaptive divergence, which must begin within the ancestral (A) or descendant (B) tumor. The new and ancestral clones are shown in dark red and pink, respectively. Tumors in HCC-4 and their genealogy are shown in panel C. The largest tumor, T1, shows genetic diversity. Since a sample from T1 (T1-3) is genetically identical with two smaller tumors, T3 and T4, the intra-tumor diversity in T1 evolves as portrayed in Fig. 3B. (D-E) Within-tumor diversity that is not part of the inter-tumor divergence. In HCC-6, the larger T1 shows intra-tumor diversity among the 5 samples. The T1-1 sample represents a distinct clone with 10 extra mutations and occupies a smaller area than the older clone as expected by the neutral model. In panel E, from each of the two larger tumors of HCC-7, three samples were taken. Within the T2 tumor, the younger clone T2-1 is smaller than the older ones. For T1, the younger T1-3 clone is also smaller than the older T1-1/T1-2.

In HCC-4 of Fig. 5C, all samples share 58 nonsynonymous substitutions. A single sample from the largest T1 tumor (T1-3) has the ancestral genotype but the majority of samples from the same tumor have 7 additional mutations, which likely drive the rapid growth of this large tumor. The pattern follows the model of Fig. 5B. In two other cases, HCC-3 (Fig. S2) and HCC-1 (Fig. S1), the same pattern of Fig. 5B is observed. In our study, the genetic variation within the old tumor does not fuel the adaptation to the new environment, which evolves *in situ* in the new tumor.

#### Neutral growth model and the diversity within tumors

The remaining cases represent the more common snapshots of intra-tumor diversity unconnected to inter-tumor divergence. In HCC-6 (Fig. 5D), the two tumors are independently derived and are “two cases in one”. Only one of these “cases” (T1) is used. This particular HCC-6 tumor is genetically diverse, with 4 of the 5 samples coming from the same clone and the last sample (T1-1) from a second clone. The latter has additional mutations and is younger, as indicated by its much smaller size. A more complicated case is HCC-7, which has three tumors, two of which were sampled. Within T1, the younger T1-3 clone is smaller than the older T1-1/T1-2 clone. Similarly, within T2, the younger T2-1 clones is smaller than the older T2-2/T2-3 clone. In contrast, between tumors, T2 is much younger than T1 but is not smaller in size (Fig. 5E). The pattern suggests selection between, but not within, tumors. In HCC-7, we have also observed a selective eve Ω clone and the two descendant T1 clones. Since we do not know the location of Ω, this could be a between-or within-tumor event. HCC-12 also shows the pattern expected of neutral evolution, with the younger clones being smaller (see Fig. S4). The fine dissection of the HCC-15 tumor offers further support of the interpretation that within-tumor diversity is often neutral (Ling et al. 2015).

## Discussion

Genetic diversity in the entire tumor(s) in an individual is a key factor in the tumors’ responses to challenges and interventions. The level of diversity is in turn governed by mutation, migration, drift and, possibly, selection. A main result in this study is the prevalence of selection in inter-tumor comparisons and, interestingly, its absence among intra-tumor clones. The former finding is likely the consequence of biases in detecting adaptively growing tumors that have reached a larger size. Beyond these visible tumors are a much larger number of tiny tumors (42), many of which may be growing at the neutral rate. Adaptive growth between tumors therefore adds more diversity to the neutral base.

For diversity within tumors, it remains controversial whether selection plays a significant role. In this and the accompanying study, we fail to reject the neutral model for evolution within tumors. Indeed, the signature of adaptive evolution is often a reduction in diversity due to selective sweeps (41, 42); the high level of observed diversity within tumors appears to contradict this general pattern. There are additional theoretical arguments against the efficacy and prevalence of selection within tumors. The arguments are generally in the form of the Hill-Robertson effect (43, 44), which reduces the effectiveness of selection in populations of high mutation rate and low recombination (See SM section B4 for details). The suggestion of neutrality is in contradiction with the prevailing view of adaptive evolution within tumors (9, 15-17, 24), a view that has not been tested against the null model of neutrality. A common argument for the adaptive view is the increasingly poor rate of patient survival with greater observed diversity (9, 45). Nevertheless, higher diversity can be the consequence, rather than the cause, of rapid proliferation as the aggressive proliferation that decreases patient survival will also generate high neutral diversity in its wake.

Within a single HCC tumor, the diversity could contain mutations at all 30 million coding sites (Ling et al. 2015). The genetic diversity among multiple tumors would be even more complex as it is the sum total of the diversities of many discrete tumors. When the cellular environment is changed by medical interventions, some previously neutral mutations may have adaptive effects on the cells, including drug resistance (the Dykhuizen-Hartl effect; see Ling et al. 2015). The total genetic diversity therefore has important implications for treatment strategies. If the diversity consists of only hundreds of coding mutations, as is generally reported, the strategy aiming at eradicating all cancer cells would be reasonable because resistant mutations may indeed be absent. However, given that the level of diversity is orders of magnitude larger than reported in the literature, the presence of many resistance mutations seems likely.

Most mutations in tumors are at very low frequencies, including those conferring drug resistance, but are likely clinically relevant since there are so many of them (Ling et al. 2015). The suggestion that drug resistance would emerge from low-frequency clones has been corroborated by the genomic analyses of cancer relapses (1-3). In both leukemia (1, 2) and solid tumors (3, 12), the recurrent tumors often carry mutations undetectable in the primary tumors. These newly observed mutations, substantial in number (3), likely pre-date drug treatment because the time before relapse (< 7 months) is too brief for so many new mutations to emerge. Indeed, analysis of the original tumors by deep sequencing identified these mutations in 90% of the examined cases (2).

Given the vast number of low-frequency mutations in tumors and the many possible mechanisms of resistance (46-48), the recurrent tumors may conceivably consist of several independent clones from the original tumor. High-resolution studies of clonal structure appear to support such an interpretation of cancer recurrence (2, 3, 47). Other types of recurrences could also be interpreted to be of multi-clonal origin. For example, if the recurrent tumor consists of 10 or more independent resistance clones, then it may not show any relapse-specific mutations, as has been commonly observed (1).

The analysis of recurrences corroborates the view of pervasive low-frequency mutations in tumors. The pre-existence of resistance has been known to impact post-treatment evolution since the time of Luria and Delbruck (1943) (49). More recently, Read et al. (2011)(50) pointed out that aggressive strategies against pathogens or cancerous cells are effective only in the absence of resistance at treatment. If resistance already exists, aggressive strategies would risk speeding up the evolution of resistance. With the understanding that two sub-populations of cells commonly co-exist, treatment-sensitive (S) and treatment-resistant (R), various strategies for administering drugs have been proposed. They include low dose and/or metronomic chemotherapy (6, 7, 51), an optimized dosing schedule (8) and adaptive therapy (4, 5).

Different strategies make different assumptions about the nature of the resistance. While clinical trials based on these different strategies are in progress (52), their eventual success will depend critically on the accuracy of the assumptions. This study and the companion analysis (Ling et al. 2015) provide a (cell) population genetic characterization of mutations in tumors. The results present both a challenge and an opportunity for treatment strategies. The challenge is the high diversity in tumors which suggests a very high probability of resistance mutations. More importantly, the very low frequencies of most mutations also represent an opportunity for retarding the evolution of resistance. Some of the strategies being tested reduce but do not eradicate S cells, which may then restrict the increase of the small number of R cells by out competing them for space and resources (4-7). This impediment to R cell proliferation will be most effective when the R/S ratio is extremely low (4, 5, 50). In this light, systematic investigations of the approximate number and frequency of mutations in tumors will be critical for designing treatment strategies.

## Acknowledgements

We are grateful to Steve Frank, Nelly Polyak, Carlo Maley, Robert Gatenby, Rick Durrett, Thomas Nagylaki, Tian Xu, Jianzhi (George) Zhang, Andy Clark, Xionglei He, and Yang Shen for inputs in various phases of this work. This work was supported by grants from the Chinese Academy of Sciences (KSCX1-YW-22) and National Science Foundation of China - 9113190 to X.M.L.

## Supplementary Materials

Materials and Methods

Supplementary Theories

Figs. S1 to S4

Tables S1 to S3

Supplementary References

